# Transcriptome-wide analysis of microRNA-mRNA correlations in unperturbed tissue transcriptomes identifies microRNA targeting determinants

**DOI:** 10.1101/2021.12.22.473932

**Authors:** Juan Manuel Trinidad Barnech, Rafael Sebastián Fort, Guillermo Trinidad Barnech, Beatriz Garat, María Ana Duhagon

## Abstract

MicroRNAs are small RNAs that regulate gene expression through complementary base pairing with their target mRNAs. Given the small size of the pairing region and the large number of mRNAs that each microRNA can control, identifying biologically relevant targets is difficult. Since current knowledge of target recognition and repression has mainly relied on *in vitro* studies, we sought to determine if the interrogation of gene expression data of unperturbed tissues could yield new insight into these processes. The transcriptome-wide repression of all the microRNA-mRNA canonical interaction sites (seed and 3’-supplementary regions, identified by sole base complementarity) was calculated as a normalized Spearman correlation (Z-score) between the abundance of the transcripts in the PRAD-TCGA tissues (RNA-seq and small RNA-seq data of 546 samples). Using the repression values obtained, we confirmed established properties or microRNA targeting efficacy, such as the preference for gene regions (3’UTR > CDS > 5’UTR), the correspondence between repression and seed length (6mer < 7mer < 8mer), and the contribution to the repression exerted by the 3’-supplementary pairing at nucleotides 13-16 of the microRNA. Our results suggest that the 7mer-A1 seed could be more repressive than the 7mer-m8, while they have similar efficacy when they interact using the 3’-supplementary pairing. The 6mer+suppl sites yielded a normalized Z-score of repression similar to the sole 7mer-A1 seeds, alerting its potential biological relevance. We then used the approach to further characterize the 3’-supplementary pairing using 39 microRNAs that hold repressive 3’-supplementary interactions. The analysis of the bridge between seed and 3’-supplementary pairing sites confirmed the optimum +1 offset previously evidenced, but higher offsets appear to have similar repressive strength. The selected microRNAs show a low GC content at positions 13-16 and base preferences that allow a sequence motif identification. Our study demonstrates that transcriptome-wide analysis of microRNA-mRNA correlations in large, matched RNA-seq and small-RNA-seq data can uncover hints of microRNA targeting determinants operating in the *in vivo* unperturbed set. Finally, we provide a bioinformatic tool to identify microRNA-mRNA candidate interactions based on sequence complementarity of the seed and 3’-supplementary regions.

## Introduction

MicroRNAs are small, single-stranded RNAs that shape the expression of messenger RNAs (mRNAs) and are essential in metazoans. Because of this, microRNAs influence all developmental processes and diseases, including cancer (Bartel, 2018). In microRNA targeting, the seed region (positions 2–7 or 2–8 of the microRNA) is the primary determinant of targeting efficacy and specificity, interacting with the 3’ untranslated region (UTR) or the coding sequence (CDS) of target mRNAs (Grimson et al., 2007; Hafner et al., 2010; Helwak et al., 2013; Moore et al., 2015; Schnall-Levin et al., 2010). Since the seed region is only composed by 6–7 contiguous nucleotides, potential target sites are frequent in the transcriptome, but not all are functionally associated with microRNA-mediated repression (Baek et al., 2008; Selbach et al., 2008; Yue et al., 2009). Although seminal studies have discovered the main rules governing microRNA-mRNA interaction, several aspects still remain unsolved and the identification of biological relevant interactions is difficult (Bartel, 2018).

It has been demonstrated that nucleotides outside of the seed of the microRNA can contribute to target recognition through complementary base pairing (Broughton et al., 2016; Duan et al., 2022; Grimson et al., 2007; Helwak et al., 2013; Moore et al., 2015; Salomon et al., 2015; Sheu-Gruttadauria et al., 2019; Wahlquist et al., 2014; Wee et al., 2012). The systematic analysis of microRNA pairing likelihood revealed the conservation of nucleotides 13–16 (Grimson et al., 2007). However, this 3’-supplementary pairing has shown little influence on on-site affinity and efficacy (Grimson et al., 2007; Salomon et al., 2015; Wee et al., 2012), and actually, after purifying selection, only about 5% of the seed-matched regulatory sites appear to involve it (Friedman et al., 2009). More recent work has resolved the crystallography of human Ago2 with microRNA-122, uncovering microRNA-mRNA interactions through the 2-8 seed and the 3’-supplementary region (nucleotides 13-16) of the microRNA (Sheu-Gruttadauria et al., 2019). They also found that perfect 3’-supplementary interactions can enhance target affinity. Additionally, they demonstrate that the seed and the 3’-supplementary regions can be connected by an unstructured loop of 1–15 nucleotides in the mRNA target. Even when the loop region has base-pairing potential, Ago2 holds a stable conformation that avoids it (Sheu-Gruttadauria et al., 2019). Further evidence indicate the benefit of microRNA 3′-pairing varies depending on the microRNA sequence and the 3′-pairing architecture, thus understanding what features determine the use of 3′-pairing was suggested to require large measurements of multiple microRNA sequences (McGeary et al., 2022). An explanation of the contradictory findings about the relevance of the 3’-supplementary interaction suggested that the differences found in the repression level between the global analyzes and those of specific microRNAs might be due to a “dilution” effect of the few strong among the most frequent weak interactions (Xiao & Macrae, 2020).

In the present study, we evaluate the contribution of the microRNA canonical seed types and the 3’-supplementary region to target mRNA repression by analyzing all the evolutionary conserved microRNAs (Agarwal et al., 2015) and all mRNA transcripts expressed in prostate tissue samples of the PRAD-TCGA cohort. MicroRNA-mRNA interaction were identified by sole base complementarity and the repressive strength was evaluated by the Spearman correlation coefficient of the expression of both interaction partners obtained from the RNA-seq transcriptomes of the PRAD-TCGA. Overall, our investigation suggests that the comparative analysis of RNA-seq transcriptomes of mRNAs and small RNAs of a large number of unperturbed tissues allows the discovery of microRNA-mRNA interactions determinants.

## Methods and Data Sets

### MicroRNAs and mRNAs transcriptomic data

RNAseq and small RNA-seq data of the TCGA-PRAD (normal and tumor conditions) were obtained from Firebrowser (firebrowser.org). The microRNA conservation scores were obtained from TargetScan 7.2 database (Agarwal et al., 2015), and the 221 microRNAs with the highest “Family Conservation score” (score 2 defined by TargetScan), hereafter denominated “conserved microRNAs”, were selected for this study.

Only microRNAs and mRNAs detected in at least 80% of the samples were included in the analysis (Supplementary Figure 1, Supplementary Table 1, see complete pipeline analysis in Figure 1).

**Figure 1.**
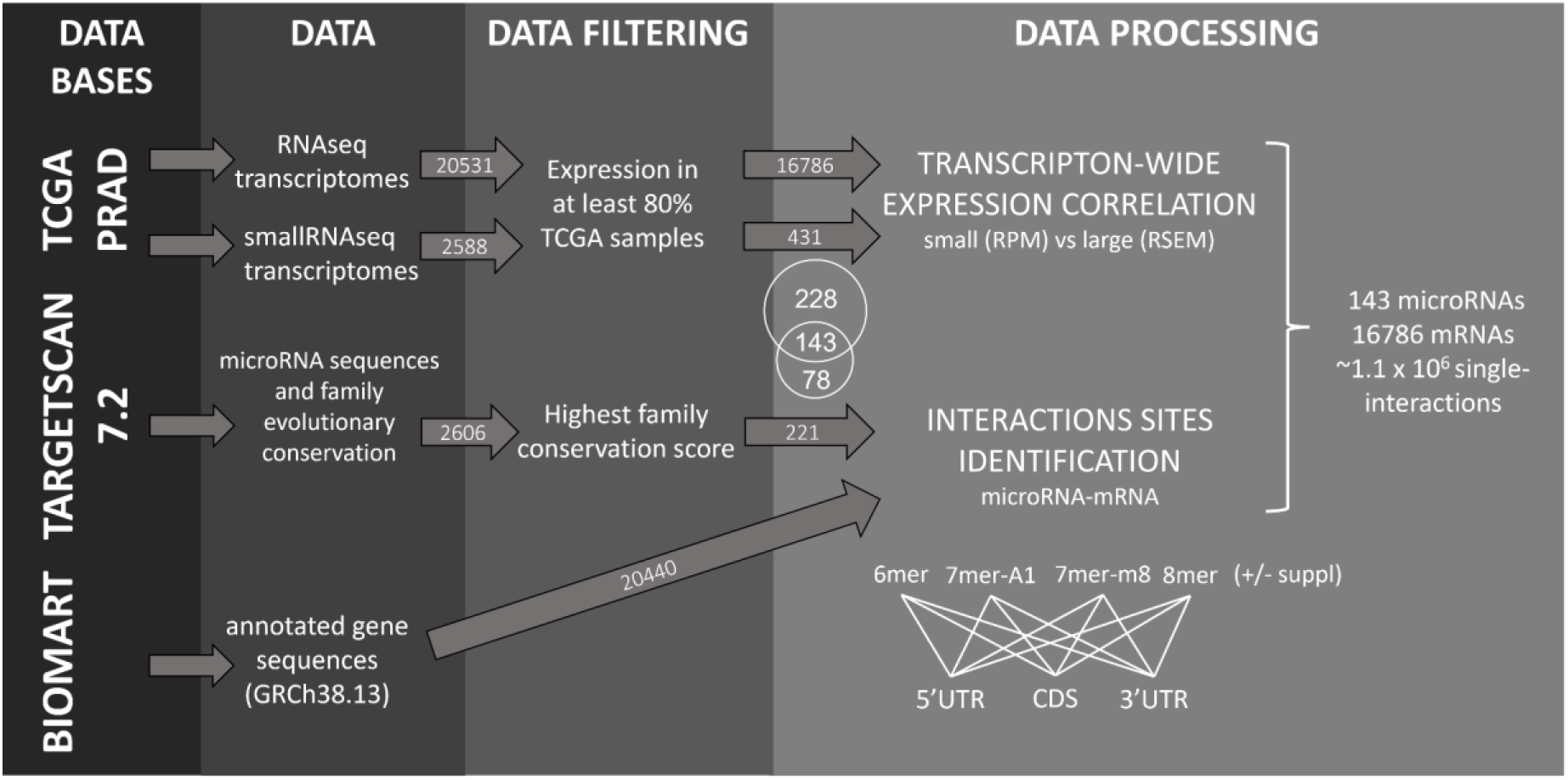
Data analysis pipeline.

### Canonical seeds and 3’-supplementary microRNA-mRNA interactions identification

Mature human microRNA sequences were obtained from TargetScan 7.2 database (Agarwal et al., 2015). Human mRNA sequences of all protein-coding genes were obtained using BioMart (Ensembl GRCh38.13).

An *in-house* script written in Python programming language was developed to identify all the sequences complementary to all the canonical microRNA seeds (as described by TargetScan) of the 143 microRNAs (script available on <u>GitHub</u>). The TargetScan nomenclature was used to name the microRNA-mRNA interaction types, which comprise 6mer, 7mer-A1, 7mer-m8, and 8mer microRNA seeds (Figure 2A). For this study, the 3’-supplementary region of an mRNA-microRNA interaction refers to microRNA nucleotides at positions 13-16 (Grimson et al., 2007; Sheu-Gruttadauria et al., 2019) (Figure 2A). For both the sole seed and seed plus 3’-supplementary region, only perfect complementary interactions with the mRNA were computed. A variable-length bridge of unpaired nucleotides can separate these two sequence modules. We investigated mRNA loops of a maximum of 15 nucleotides length based on previous functional studies (Sheu-Gruttadauria et al., 2019). The difference in the length of the microRNA and the mRNA unpaired bridges is referred to as the “offset” of the interaction involving 3’-supplementary sites.

**Figure 2.**
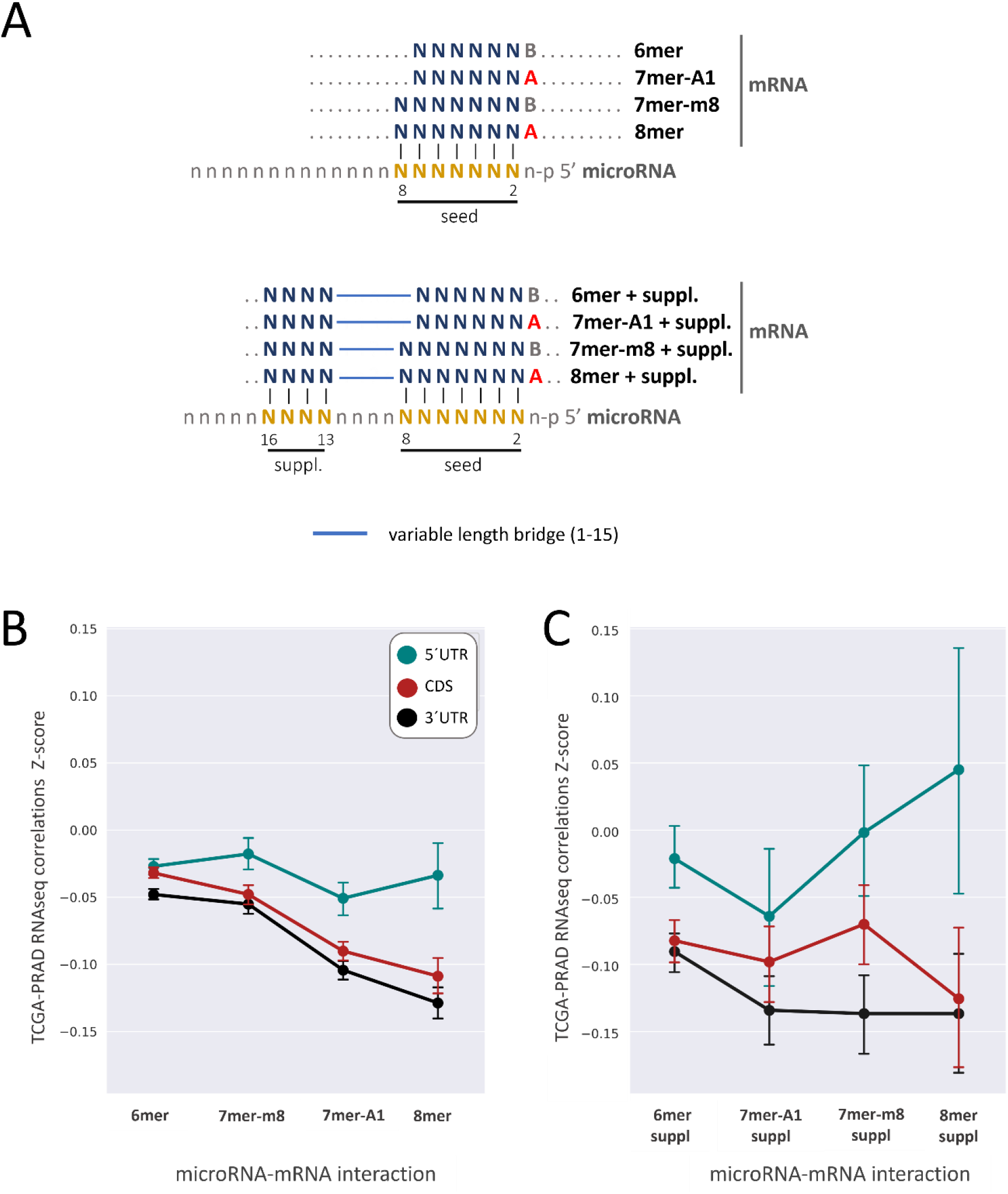
Repression of mRNA-target interactions with different pairing sites. The average Z-score correlations between the 143 conserved microRNAs selected and their predicted target mRNAs are represented for microRNA/mRNA interactions at the CDS, 5’UTR, and 3’UTR. Vertical bars represent the confidence interval (95%). Letter A in the sequences corresponds to Adenine, and Letter B corresponds to the other three nucleotides except Adenine.

We identified the eight types of putative microRNA-mRNA interactions described using Python regular expressions, comprising 6mer, 7mer-A1, 7mer-m8, 8mer, 6mer+suppl 7mer-A1+suppl, 7mer-m8+suppl, 8mer+suppl (Figure 2A). We chose the transcript variant with the highest number of putative microRNA interactions for each gene. In addition, per each microRNA, we selected the mRNAs that presented only one predicted microRNA-mRNA interaction, pondering the CDS, 3’UTR and 5’UTR independently. As a result, the computed mRNAs only have a single microRNA site at the region considered for every given microRNA analyzed, thus avoiding the co-occurrence of the sites.

### MicroRNA-mRNA pairs correlation analysis

For the selected list of microRNA and mRNAs, the normalized Spearman correlation coefficient (Z-score centered on microRNA) for the abundance of all putative microRNA-mRNA interacting pairs were calculated with the Python’s Pandas library (McKinney Wes, 2010) using TCGA RNA-seq (mRNAs) and small RNA-seq (microRNAs) normalized data (546 tissue samples) obtained from FireBrowse (firebrowse.org).

### Statistical analysis

Unless specified, statistical differences were assessed by Two-tailed T-test, and the p-values were adjusted by Bonferroni correction, using SciPy library version 1.6.2. Fisher exact test was performed using SciPy library version 1.6.2. In all cases, adjusted p-values < 0.05 were considered significant. All the plots were generated using Seaborn library 0.11.2.

### Executable files

The executable files to search for microRNA/mRNA interaction sites were developed using PySimpleGUI 4.55.1 (graphical user interface) and PyInstaller 4.10 and are available on <u>GitHub</u>.

## Results

### Identification of microRNA-mRNA interaction sites in the PRAD-TCGA transcriptome based on nucleotide base complementarity

Given the tissue specificity of microRNA expression, we performed this study within a single tissue type. We chose the PRAD cohort because it is large and extensively studied for microRNAs (Lin et al., 2021; Wei et al., 2020; Yang et al., 2019; Ye et al., 2018); in addition, PRAD is a very heterogeneous cancer thus biases result in of cancer subtypes are unlikely (Abeshouse et al., 2015). We investigated the expression of the conserved human microRNAs (221) and all the human protein-coding transcripts annotated in Ensembl (20,440) in the 546 samples of the PRAD-TCGA cohort (20,531) (see pipeline analysis in Figure 1). To increase the signal/noise ratio, we filtered out low expressed transcripts, arbitrarily defined as those undetected in ≥ 20% of the samples, narrowing the list of microRNAs and mRNA to 143 and 16,786, respectively (Supplementary Table 1 and Supplementary Figure 1). Aiming to compare the repressive power of different microRNA pairing sites, we studied 6mer, 7mer-A1, 7mer-m8, and 8mer microRNA seeds with or without the 3’-supplementary pairing regions, defined as the four nucleotides at positions 13-16 of the microRNA (hereafter denominated 6mer+suppl, 7mer-A1+suppl, 7mer-m8+suppl, and 8mer+suppl) (Figure 2A). The choice of the 3’-supplementary pairing site was based on the literature since mounting evidence of its repressive contribution has been published (references in Bartel 2018, Sheu-Gruttadaria et al., 2019). We determined all the mRNA sites with perfect complementarity to these eight canonical microRNA pairing sites, presuming they represent putative microRNA binding sites. We performed this analysis independently for each of the three regions of the mRNAs (5’UTR, CDS, and 3’UTR). The examination of seed sequence complementarity between the 143 microRNAs and the 16,786 mRNAs identified 3,065,731 putative microRNA interaction sites on the mRNAs (285,237 at 5’UTR, 1,226,568 at CDS, 1,553,926 at 3’UTR). Since each mRNA can bear more than one putative microRNA site for a given microRNA, in subsequent calculations, we only included the interactions involving microRNAs with a unique site in the mRNAs. Therefore, if an mRNA has more than one site for a given microRNA, these interactions were excluded from the analysis. Yet, if the same mRNA had sites for other microRNAs, these interactions were compiled. We performed this calculation for each of the three mRNA regions independently. We iterated this analysis for each of the 143 selected microRNAs, obtaining a total of 1,116,787 sites (193,203 at 5’UTR, 443,323 at CDS, 480,261 at 3’UTR), corresponding to an average microRNA interaction number of 1,351±621 at the 5’UTR, 3,100±748 CDS, and 3,358±602 at the 3’UTR (Supplementary Table 2). As expected, for the eight interaction types in the three regions studied, the number of predicted interactions decreases with the seed size and the incorporation of the 3’-supplementary regions (Supplementary Figure 2).

### Inference of microRNA activity from microRNA-mRNA expression correlation in the PRAD-TCGA transcriptomes

The availability of matched RNAseq and small-RNAseq and transcriptomes of hundreds of tissues due to global cancer initiatives allows the high-throughput transcriptome-wide assessment of microRNA-mRNA correlations. Our approach evaluates the strength of microRNA-mRNA interaction by the Spearman correlation coefficient of the expression of both molecules calculated from the RNAseq data of the 546 patient tissues (normal and tumor conditions) of the PRAD-TCGA cohort (Figure 1). Since the microRNAs can exert different effects on the expression of their target genes, we included both positive and negative correlations between the mRNA-microRNA pairs. Additionally, in the absence of consensus about the threshold value of significant biological correlation expected in tissues for direct microRNA repression, we analyzed all the correlations regardless of their magnitude.

### Validation of microRNA repressive mechanisms using microRNA activity inferred from microRNA-mRNA expression correlations in PRAD-TCGA

We initially sought to evaluate if the sole correlation of microRNA-mRNA pairs abundance in the tissues validates established microRNA repression rules. The comparison of a Z-score of microRNA-mRNA correlation for each type of seed showed that repression levels increase with seed’s length, i.e., the number of nucleotide bases involved in the interaction (6mer < 7mer-m8 < 7mer-A1 < 8mer) (Figure 2B and Supplementary Figure 3) (Table 1 and Supplementary Table 3), thus confirming a previously established rule of microRNA targeting (Friedman et al., 2009; Grimson et al., 2007; Hafner et al., 2010). These Z-score differences are statistically more significant for intra 3’UTR (average p-value = E-25), followed by intra CDS (average p-value = E-16) seed comparisons. Meanwhile, only 6mer/7mer-A1 and 7mer-A1/7mer-m8 differences are marginally significant at the 5’UTR (p-value 1.14 E-02 and 2.93 E-03, respectively). As expected, the strength of the mRNA target repression inferred from the correlations increases with the length of the microRNA/mRNA pairing region (Figures 2B and 2C), although the significance of the differences is influenced by the number of interactions in each dataset. Thus, the hierarchy of site efficacy is as follows: 8mer > 7mer > 6mer (Grimson et al., 2007; McGeary et al., 2019; Nielsen et al., 2007) (Figure 2B). In contrast to the previous findings (Friedman et al., 2009; Grimson et al., 2007), the Z-score indicates that the 7mer-A1 seed interaction is significantly more repressive than the 7mer-m8 in the three regions of the mRNA studied (Figure 2B). Additionally, the 7mer-m8 shows no difference from the 6mer seed in the UTRs (Figure 2B).

**Table 1.**

| Group1 | Group2 | *p-value* |  |  |
| --- | --- | --- | --- | --- |
|  |  | 5'UTR | CDS | 3'UTR |
| 6mer | 7mer-m8 | n.s. | 6.09E-03 | n.s. |
| 6mer | 7mer-A1 | 1.14E-02 | 2.36E-42 | 4.82E-45 |
| 6mer | 8mer | n.s. | 1.74E-25 | 9.23E-35 |
| 7mer-m8 | 8mer | n.s. | 2.24E-13 | 4.41E-23 |
| 7mer-A1 | 7mer-m8 | 2.93E-03 | 3.94E-14 | 9.48E-21 |
| 7mer-A1 | 8mer | n.s. | n.s. | 1.24E-02 |
| 6mer | 6mer suppl | n.s. | 2.84E-08 | 9.90E-07 |
| 6mer | 7mer-m8 suppl | n.s. | n.s. | 3.66E-08 |
| 6mer | 7mer-A1 suppl | n.s. | 1.71E-04 | 2.48E-09 |
| 6mer | 8mer suppl | n.s. | 1.23E-02 | 3.72E-03 |
| 7mer-m8 | 7mer-m8 suppl | n.s. | n.s. | 1.74E-06 |
| 7mer-m8 | 8mer suppl | n.s. | n.s. | 1.52E-02 |
| 6mer suppl | 7mer-m8 | n.s. | 2.84E-03 | 6.92E-04 |
| 6mer suppl | 8mer | n.s. | n.s. | 1.74E-03 |
| 7mer-A1 suppl | 7mer-m8 | n.s. | 2.20E-02 | 2.43E-07 |

Figure 2C presents an identical analysis carried out for the interaction sites involving seeds with base pairing possibilities at the 3’-supplementary regions. Again, most of the comparisons show more repression for longer seeds, which is more notoriously at the CDS and 3’UTR; the comparatively lower significance of the observed differences could be influenced by the smaller number of interactions considered in comparison to the sole seeds analysis (Figure 2B). The comparison of sole seeds and seeds plus 3’-supplementary regions shows that the 3’-supplementary interaction mainly contributes to repression, particularly at the 3’UTR and the CDS (Figure 2B and 2C and Supplementary Figure 3), although only the 6mer (CDS and 3’UTR) and 7mer-m8 (3’UTR) differences are statistically significant (p-value < 0.05) (Table 1). This finding from tissue data supports the contribution of the 3’-supplementary 13-16 nt microRNA-mRNA interaction to the microRNA repression that has been proposed using *in vitro* approaches (Grimson et al., 2007; Helwak et al., 2013; Moore et al., 2015; Sheu-Gruttadauria et al., 2019; Xiao & Macrae, 2020). Additional validation of established rules of microRNA targeting mechanism comes from the observation that the relative magnitude of the repression of the microRNA/mRNA interactions is 3’UTR > CDS > 5’UTR (Figure 2B-C) (Grimson et al., 2007; Hafner et al., 2010; Helwak et al., 2013), which is supported by the p-values of the comparison among the sites in the three different regions (Table 2). This finding provides another proof of principle for using microRNA/mRNA correlations in unperturbed tissues to study *bona fide* microRNA repression characteristics.

**Table 2.**

| Group1 | Group2 | *p-value* |  |  |  |
| --- | --- | --- | --- | --- | --- |
|  |  | 6mer | 7mer-A1 | 7mer-m8 | 8mer |
| 3UTR | 5UTR | 5.98E-09 | 2.69E-14 | 4.34E-07 | 7.30E-12 |
| 3UTR | CDS | 2.62E-08 | 1.37E-02 | n.s. | n.s. |
| 5UTR | CDS | n.s. | 6.25E-08 | 6.82E-05 | 2.06E-07 |
| Group1 | Group2 | *p-value* |  |  |  |
|  |  | 6mer suppl | 7mer-A1 suppl | 7mer-m8 suppl | 8mer suppl |
| 3UTR | 5UTR | 2.81E-06 | 3.62E-02 | 1.14E-05 | 1.29E-03 |
| 3UTR | CDS | n.s. | n.s. | 5.11E-03 | n.s. |
| 5UTR | CDS | 7.30E-05 | n.s. | n.s. | 4.49E-03 |

### Insight into the characteristics of microRNA-mRNA interaction involving beneficial 3’-supplementary pairing regions withdrawn from microRNA/mRNA correlations

Our previous results drew our attention to the relevance of the 3’-supplementary region interaction for all types of seed; thus, we sought to investigate if the analysis of the microRNA activity inferred from the tissue transcriptomes could uncover features of the microRNA/mRNA pairing involving 3’-supplementary regions. For the following analyzes, we focus on the seed plus 3’-supplementary sites of 6mer+suppl (at 3’UTR and CDS) and 7mer+suppl. (at 3’UTR), since they showed a statistically significant increase in target repression compared to sole seeds (Table 1); therein, we selected the microRNAs that exert more repression on targets bearing 3’-supplementary interactions than on targets without 3’-supplementary pairing possibilities, as determined by a T-test of Z-score differences (p-value ≤ 0.05) (Supplementary Table 4). Overall, we found 39 repressive microRNAs in one or more of the three gene regions studied and 18 microRNAs with the opposite behavior (i.e., positively correlated with their target mRNAs). If each dataset is analyzed separately, the microRNAs that follow or oppose the repressive behavior are comparatively 19 vs. 11 (6mer 3’UTR, p-value = 0.2 binomial test of significance), 16 vs. 4 (7mer-m8 3’UTR, p-value = 0.01 binomial test of significance), and 21 vs. 8 (6mer CDS, p-value = 0.02 binomial test of significance). The 39 repressive microRNAs have no significant difference in expression (Supplementary Figure 4) or number of mRNA targets (Supplementary Figure 5) than the total 143 microRNAs analyzed (p-value ≤ 0.05).

High-throughput approaches showed that nucleotides within positions 8-13 of the microRNA are not paired in the human AGO-microRNA-target complex (Hafner et al., 2010). Consistently, it was later shown that the seed and the 3’-supplementary regions could be bridged by an unstructured target loop of 1-15 nucleotides when complexed with AGO2, even when central complementary bases are available for pairing (Sheu-Gruttadauria et al., 2019). Moreover, 8mer+suppl interactions with bridges up to 10 nucleotides long in the target mRNA were more repressive than 8mer sole seeds. The same study proposes that interactions involving 3’-supplementary regions with high GC content can be established using bridges of up to 15 nt (Sheu-Gruttadauria et al., 2019). Since the bridge between the seed and the 3’-supplementary region of the microRNA is five nucleotides for the 6mer+suppl (8-12 nt) and four nucleotides for 7mer-m8+suppl (9-12 nt), the same length at the mRNA bridge means no loop formation in any of the two molecules (zero offset) (Figure 3A). Longer and shorter mRNA bridges imply a loop formation in the mRNA (positive offset values) or microRNA (negative offset values).

**Figure 3.**
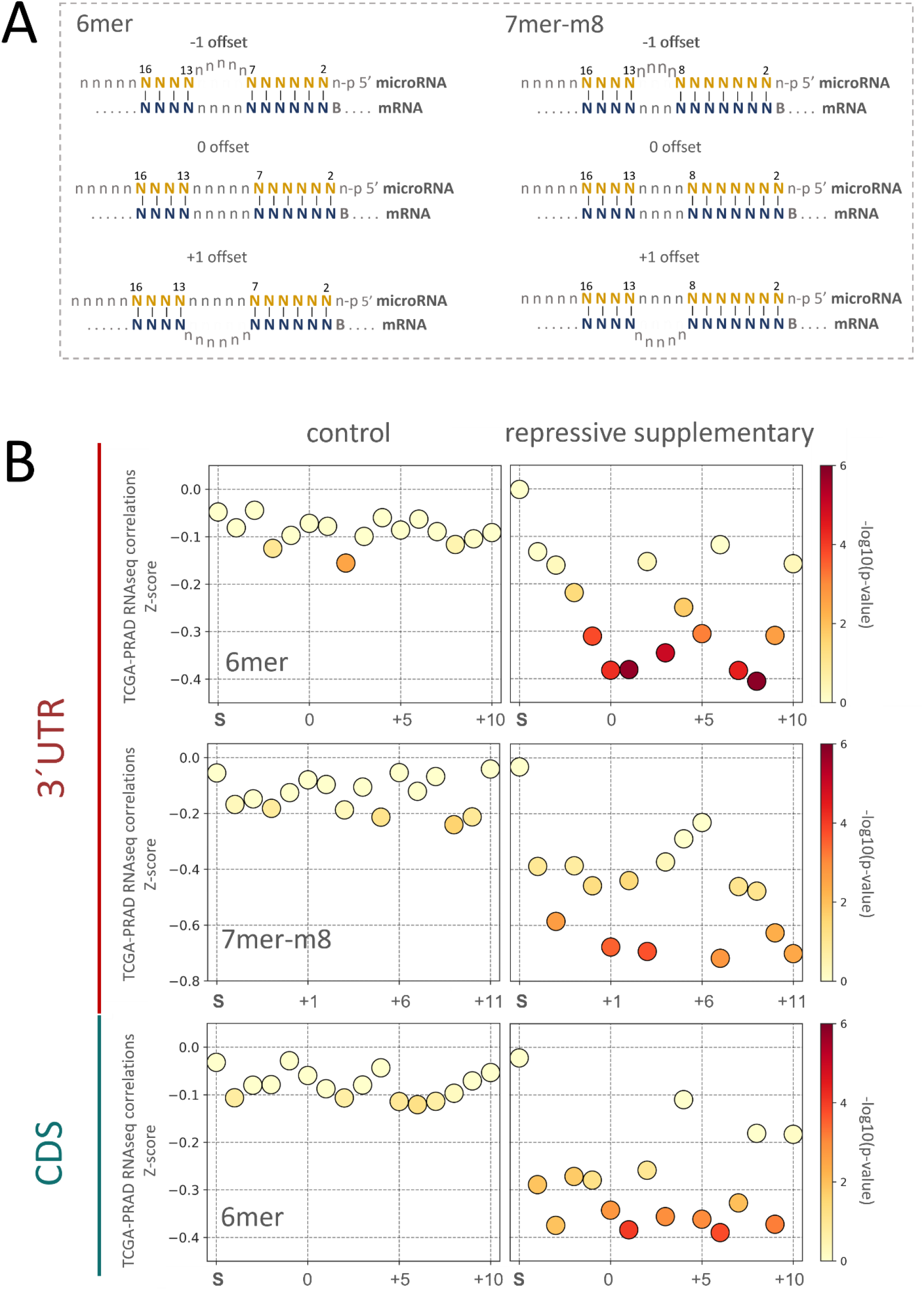
Contribution of the mRNA bridge length to the microRNA repression. **A)** Graphic representation. The bridge region is defined as the unpaired nucleotides from the end of the seed to the beginning 3’-supplementary region. Since microRNA/mRNA 3’-supplementary interaction occurs at positions 13-16 of the microRNA, 6mer and 7mer-m8 seeds have 5 and 4 nt microRNA bridges, respectively; thus, these mRNA bridge lengths mean no loop formation for the respective seeds and are denominated “zero offset”. Positive offset values imply loop formation in the mRNA and negative offset values in the microRNA. **B)** Analysis of bridge length effect on mRNA target repression (Z-transformed score) using the TCGA transcriptomes and the 143 total microRNAs included in this study as a control (left panels) and the 39 microRNAs with repressive 3’-supplementary interactions (right panels). The region of the genes and the type of seeds analyzed are indicated. The -log10(p-values) of the differences between each seed+suppl length and the respective sole seed interactions are indicated by the color intensity bar. Letter S stands for “seed” and indicates the Z-score of sole seed interactions.

Seeking to assess if the Z-score of microRNA repression withdrawn from the tissue PRAD-TCGA data was influenced by the offset of the microRNA/mRNA interaction, we analyzed the repression exerted by the 39 repressive microRNAs using the different offsets defined for their predicted interaction with their mRNA targets. As a control, we performed the same analysis for all the interactions involving 3’-supplementary regions pairing of the 143 microRNAs. As expected, their Z-scores are higher than those of the 39 repressive microRNAs, regardless of the seed type (6mer or 7merm8), gene location (CDS or 3’UTR), or the offset of the interaction (Figure 3B). Meanwhile, the 3’-supplementary interactions of the 39 repressive microRNAs show heterogeneous repression depending on the offset (Figure 3B, right panels). For the 6mer interactions occurring at the 3’UTR, a gradual gain of repression is observed with shorter microRNA loops (negative offsets), reaching their maximum repression at 0 offset. On the contrary, the length of the mRNA loops (offsets +1 to +10) is not directly proportional to the repressiveness of the 6mer interaction at the 3’UTR, yet the +1 offset is one of the most repressive lengths observed. A similar pattern of repression is depicted for the two other datasets (6mer CDS and 7mer-m8 3’UTR), though the differences are smoother, probably due to the smaller number of interactions composing them. To add up the information from the three datasets, we scaled their Z-scores using a minimum-maximum scaling (to normalize for their different Z-score ranges), and then we combined the three values. This integrated analysis indicates that the length of the microRNA loop is inversely proportional to the repression, while a one nucleotide loop in the mRNA (+1 offset) provides maximum repression (Supplementary Figure 6). Consistently, the optimum repression has been shown at +1 (McGeary et al., 2022; Sheu-Gruttadauria et al., 2019). Nevertheless, our results suggest that other offsets may produce similar repression.

Since base content bias has been previously proposed at the 3’-supplementary microRNA interaction sites (Sheu-Gruttadauria et al., 2019), we analyzed the GC content of the microRNAs with repressive 3’-supplementary pairing in comparison with the complete set of microRNAs. We observed a notorious decrease of the GC content in the stretch between positions 13-16 of the microRNAs with a repressive supplement for the three types of interactions analyzed (Figure 4A). In addition, positions 5 and 19 of 6mer 3’UTR sites have a GC content deviated from the total. The base composition analysis per position shows a preference for base A at positions 13-15 of the 3’-supplementary pairing region of the microRNA and for base T at position 16 for the three interaction types analyzed (Figure 4C-F). The logo represented in Figure 4B resulted from the comparison of the full-length 39 microRNA mature sequences with the 143 microRNAs obtained using the Discriminative Mode of the Motif Discovery of The MEME Suite (Bailey et al., 2015); it is supported by 16 microRNAs (E-value = 4.7e-15). Although the enrichment of base T and A observed at positions 1 and 2 of all the microRNA (Figure 4C) has been previously described in mammals and other species (Friedman et al., 2009; B. Wang, 2013), the logo enriched in the 39 selected microRNAs suggests an overrepresentation of these bases (Figure 4B).

**Figure 4.**
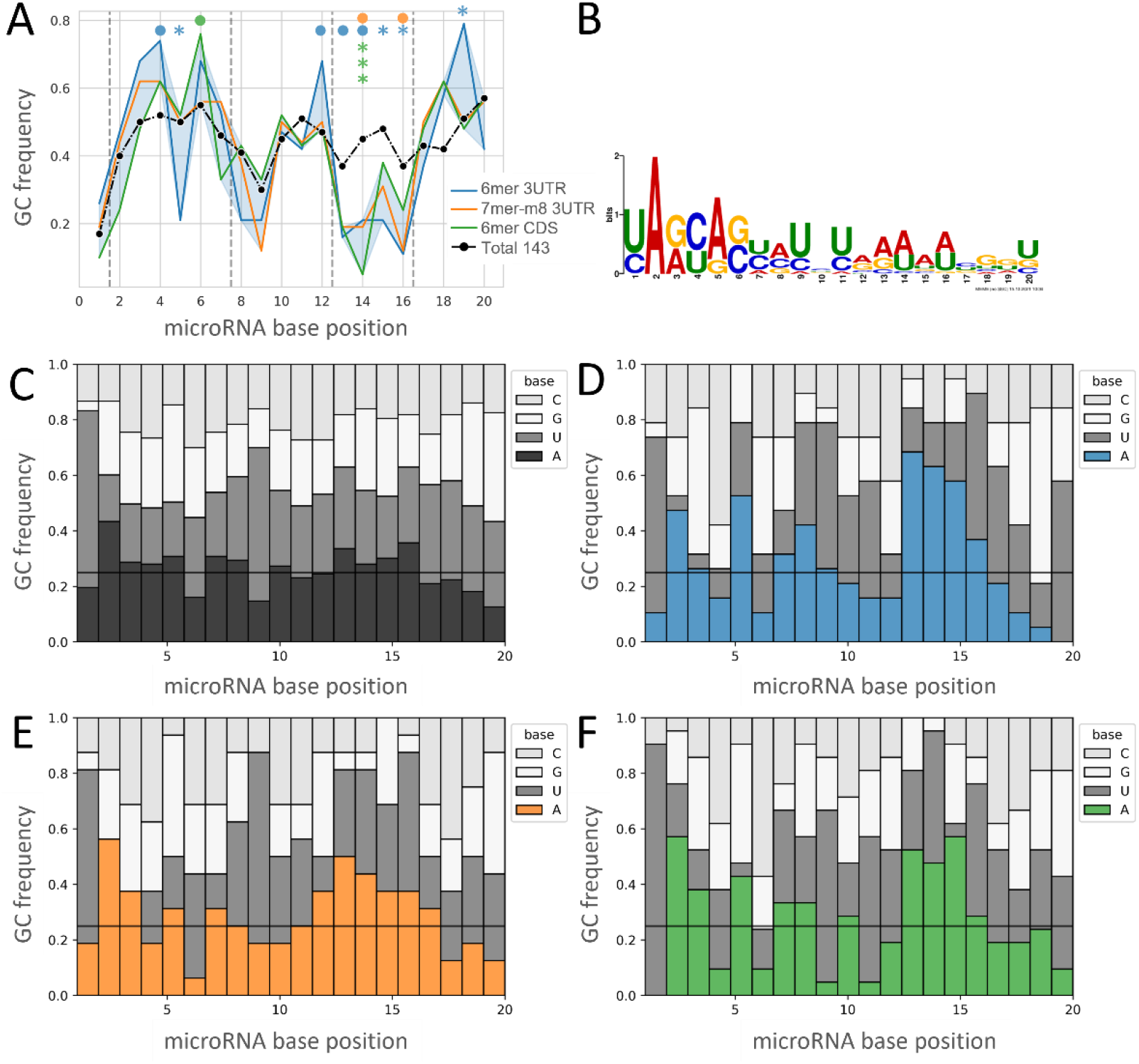
Nucleotide composition of the microRNAs with repressive 3’-supplementary interactions. **A)** Distribution of GC content along the positions of the microRNAs with the three repressive 3’-supplementary interactions indicated (colored) versus the whole set of microRNAs analyzed (black). Fisher exact test was performed to compare each of the represented sites with the total 143 microRNAs (● < 0.1, * < 0.05, *** < 0.001). **B)** MicroRNA site identified by the MEME suite (Discriminative Mode), using the 39 versus all the 143 microRNAs. The motif is supported by 16 of the 39 microRNAs (E-value = 4.7e-15). **C-F)** Base composition per microRNA nucleotide position. Seed types are represented by colors as indicated in A (C: all microRNAs, D: blue/6mer 3’UTR, E: orange/7mer-m8 3’UTR, and F: green/6mer CDS). Horizontal lines represent the ¼ frequency expected for a random nucleotide distribution.

Overall, microRNAs that establish repressive 3’-supplementary interactions with their targets seem to have low GC content at positions 13-16. Although the 6mer analysis suggests additional compositional skews, more interactions should be needed to confirm these observations.

### An executable tool to identify the microRNA-mRNA seed+suppl interactions

Finally, we developed two Windows compiled executable scripts to search for seed+suppl interaction in human genes solely based on mRNA/microRNA perfect base-pairing potential. One tool retrieves all the mRNA targets for a given microRNA, and the other retrieves all microRNAs that interact with a given mRNA transcript. The executable files identify 6-8mer seeds and 3’-supplementary interactions intervened by 1-15 nucleotides between the seed and the 3’-supplementary pairing regions. The search discriminates CDS, 3’ and 5’ UTR, and all annotated transcript variants. The software, manual, and files are available on <u>GitHub</u>.

## Discussion

The interaction between microRNAs and their target mRNAs can be inferred from the regulation of the target mRNA and protein levels provoked by the enforced modulation of the microRNA abundance in experimental settings (Baek et al., 2008; Grimson et al., 2007; Hafner et al., 2010; Helwak et al., 2013; Selbach et al., 2008). Despite their contribution to the discovery of target recognition features, these experimental approaches have the disadvantage of being limited to few microRNAs, which are generally assessed *in vitro* and in stablished laboratory cell lines. Meanwhile, evidence of *in vivo* microRNA regulation has been withdrawn from the correlation between microRNA/mRNA pairs abundance in unperturbed cells or tissues, which is usually used as a criterion to support the microRNA regulation on single mRNA targets (Ahmed et al., 2018; Elton & Yalowich, 2015; Goldman et al., 2020; Li et al., 2014; T. T. Wang et al., 2019). Since Cancer Genome projects involving multi-omics methods have provided matched RNAseq and small-RNAseq of hundreds of tissues, they allow the high-throughput transcriptome-wide assessment of microRNA-mRNA expression correlations in patient tissues.

Here we sought to investigate if the sole analysis of microRNA-mRNA correlations in large tissue datasets can validate established microRNA-mRNA pairing rules, which may indicate its putative power to uncover novel findings. We thus performed a transcriptome-wide approach to measure the correlation between all the mRNA transcripts and conserved microRNAs expressed in the PRAD-TCGA cohort. Our strategy is solely based on perfect Watson-Crick base complementarity at the 2-7 or 2-8 and the 13-16 positions of the microRNA seed and 3’-supplementary regions to assign candidate microRNA interaction sites on the target mRNAs. Although this sole criterion is permissive, it has the advantage of reducing potential biases of the microRNA/mRNA target prediction algorithms (target site conservation, the free energy of the interaction, site accessibility, target-site abundance, local AU content, GU wobble pairing in the seed, exact position in the transcript, length of the transcript regions, machine learning features) (Peterson et al., 2014). In support of this notion, it has been stated that a simple text search yields predictions more reliable than some common algorithms (Bartel, 2018). Furthermore, the use of this single criterion maximizes the number of interactions, thus increasing the statistical significance of the findings. Another advantage of our approach is the use of endogenous level of transcript abundance, since concerns have been raised about artifactual observations due to non-physiological stoichiometry of the perturbation-based approaches (Xiao & Macrae, 2020). Meanwhile, a limitation of the study is the exclusion of seed-site interactions with mismatches. In addition, the lack of inclusion of standard target prediction rules may decrease the biological meaning of the findings. Nevertheless, the choice to analyze conserved microRNAs favors the compliance to mechanistically conserved microRNA-target interaction properties.

Our initial analysis allowed the confirmation of well-established rules or microRNA target regulation, such as the dependence of repression on the gene region (3’UTR > CDS > 5’UTR), the seed length (6mer < 7mer < 8mer), and the contribution of the 3’-supplementary pairing at 13-16 of the microRNA (Agarwal et al., 2015; Bartel, 2009, 2018; Grimson et al., 2007; McGeary et al., 2022; Moore et al., 2015; Sheu-Gruttadauria et al., 2019). It is worth noting that, in terms of inferred repression, the 6mer+suppl interaction is not significantly different from the 7mer-A1, which raises the awareness of the possibly underestimated relevance of the 6mer interaction *in vivo*. Although the 6mer site *per se* is known to be less specific for repression than longer seeds and is classified as a “marginal site” (Agarwal et al., 2015; Grimson et al., 2007), the addition of a 3’-supplementary four nucleotide base pairing (microRNA position 13-16) can confer a more significant Argonaut mediated repressive activity. Additionally, contrary to previous reports (Grimson et al., 2007; Nielsen et al., 2007), our data suggest that the 7mer-A1 seed is significantly more repressive than the 7mer-m8 in the three transcript regions analyzed. Further investigation is needed to understand if this unexpected finding is due to the nature of the models studied (*in vitro* vs. *in vivo*), the bioinformatic pipelines used, or another source variation.

The functionality of the 3’-supplementary pairing in microRNA-mRNA interactions was described several years ago (Grimson et al., 2007), and further insight into its molecular basis was recently achieved (Bibel et al., 2022; McGeary et al., 2022; Sheu-Gruttadauria et al., 2019; Xiao & Macrae, 2020). Unlike the first studies that used a 2-6 nt bridge length (Grimson et al., 2007), recent studies showed that up to 15 nt long loops on the target mRNA could bridge the seed and the 3’-supplementary region (McGeary et al., 2022; Sheu-Gruttadauria et al., 2019). The discrepancies about the relative contribution of the 3’-supplementary pairing to the target repression among the aforementioned studies have been ascribed to the different methodological strategies of the analyses (Xiao & Macrae, 2020). In this context, we thought that the study of large number of interactions in tissue samples would contribute to the discovery of characteristics of the 3’-supplementary interaction, such as the bridge length, base preferences, or repression strength.

Our approach found that 3’-supplementary nucleotides pairing significantly enhance the repressive activity of 6mer and 7mer-m8 seeds in at least 39 conserved microRNAs in the 3’UTR and CDS of mRNAs targets. In agreement with previous reports the optimal offset for repression was identified at +1 (McGeary et al., 2022; Sheu-Gruttadauria et al., 2019). Moreover, the pattern of repression for offsets larger than +1 suggests that the 3’-supplementary pairing can be equally repressive at higher offsets, such us +7. Heterogeneous 3’ microRNA architecture may explain the heterogeneity of the repression per offset (McGeary et al., 2022). Interestingly, the two binding modes recently reported for compensatory interactions by McGeary et al. 2022 (offset 0-1 and +3-4) are supported by our findings of a second repressive offset at +3-4 offsets.

We also identified sequence patterns enriched in the 39 repressive supplementary microRNAs, including a low GC content with a higher adenine proportion at the 3’-3’-supplementary region and a preferred sequence motif, whose relevance must be validated by additional methods. Conversely, a low GC content in this region was previously associated with a lower affinity of the interaction evaluated *in vitro* using a specific microRNA-RNA target pair (McGeary et al., 2022; Sheu-Gruttadauria et al., 2019; Xiao & Macrae, 2020); this discrepancy may be due to the diversity of microRNAs studied, the context of the interaction (*in vitro*/*in vivo*) or to still unknown factors.

Our study represents a novel high-throughput approach to investigate the microRNA-mRNA target interaction by withdrawing patterns from transcriptome-wide expression correlations in large unperturbed tissue datasets. The findings confirm existing knowledge, supporting the biological meaning of the approach, and leading to novel hypothesis.

## Author Contributions

Conceptualization, M.A.D., R.S.F.; methodology, R.S.F, J.M.T.; software, R.S.F, J.M.T., G.T.; formal analysis, M.A.D., R.S.F, J.M.T.; data curation, R.S.F, J.M.T.; writing—original draft preparation, J.M.T. and M.A.D.; writing—review and editing M.A.D., R.S.F., J.M.T.; project administration, M.A.D. and B.G.; funding acquisition, M.A.D. All authors have read and agreed to the published version of the manuscript.

