## Supplementary Figures for "Transcriptome-wide analysis of microRNA-mRNA correlations in unperturbed tissue transcriptomes identifies microRNA targeting determinants"

### Supplementary material - Figures

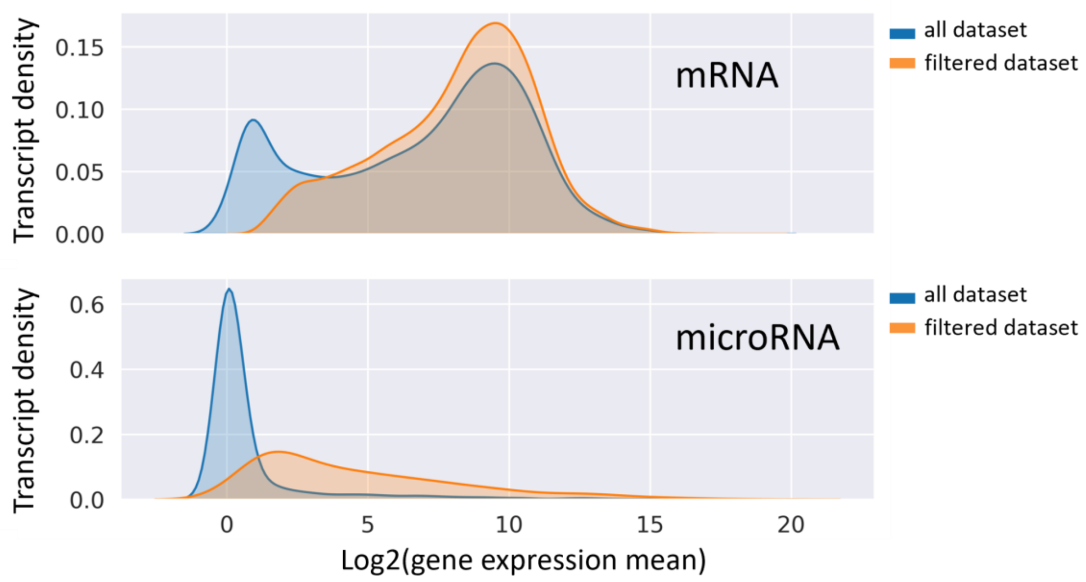


**Supplementary Figure 1.** *Density distribution for microRNA and mRNA expression on* the TCGA-PRAD dataset before and after filtering detection in 80% of the samples.

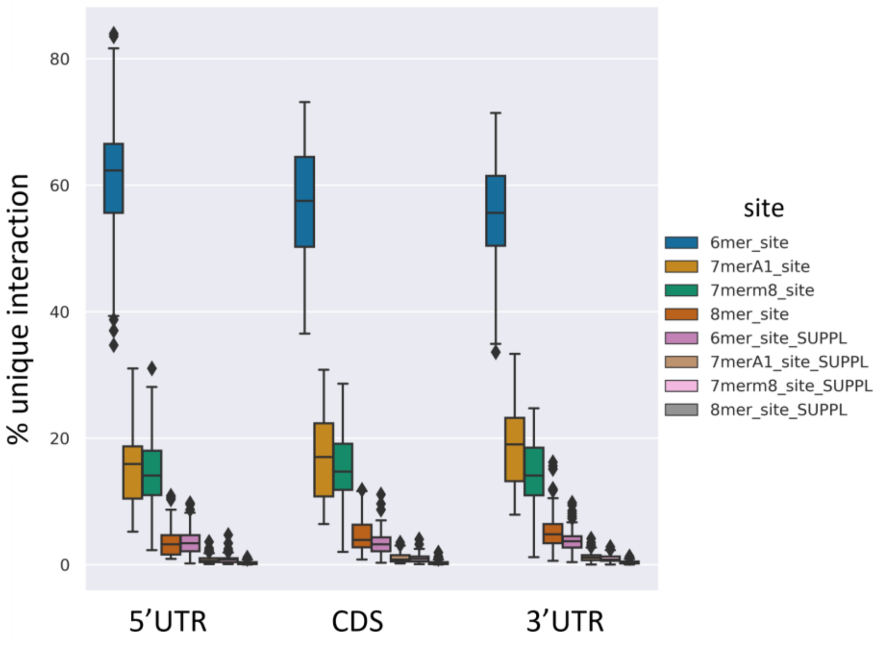


**Supplementary Figure 2.** *Percentage of unique interactions per microRNA site types by the transcript regions*, as indicated in the side-colored labels and the x-axis labels. Data calculated for the 143 filtered in microRNAs.

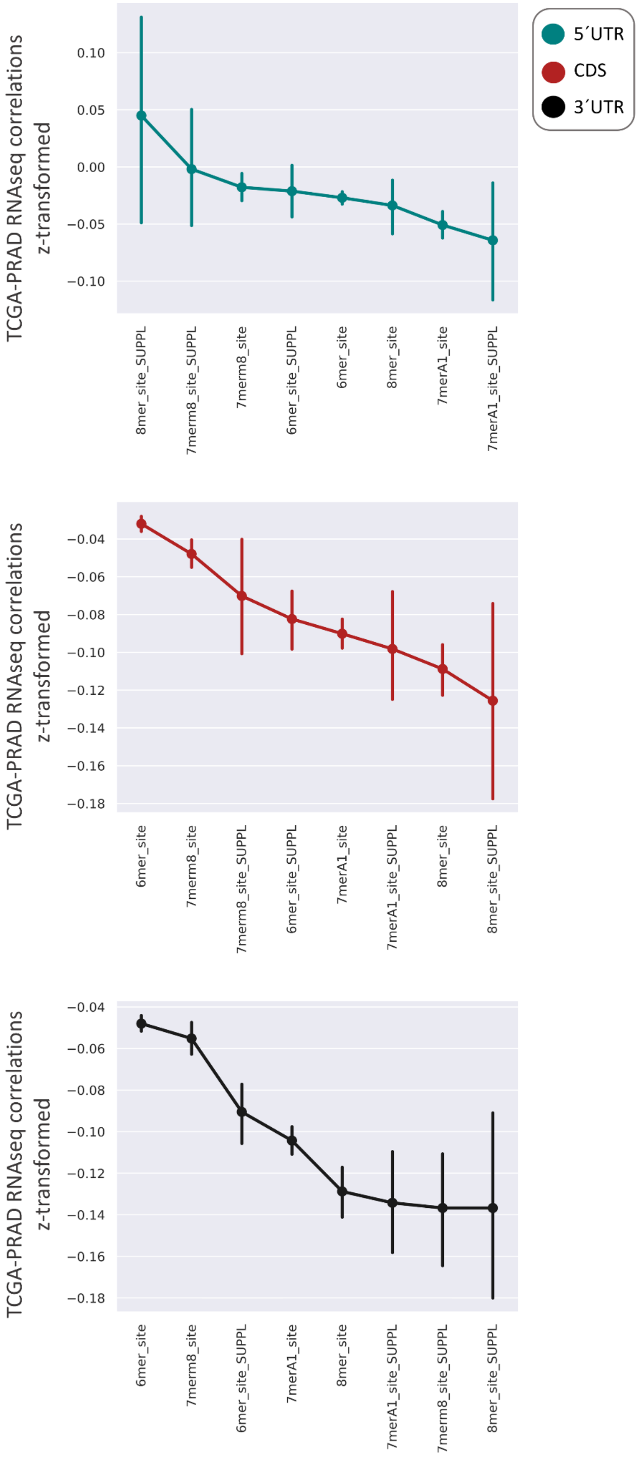


**Supplementary Figure 3.** *Repressive strength of the canonical microRNA interactions by transcript region.* Data from Figure 2 was plotted by repressive strength order. The average Z-transformed correlations between the 143 conserved microRNAs and their predicted target mRNA are represented for the coding regions (CDS) and the 5´ and 3´ untranslated regions (UTR). Vertical bars represent the confidence interval (95%).

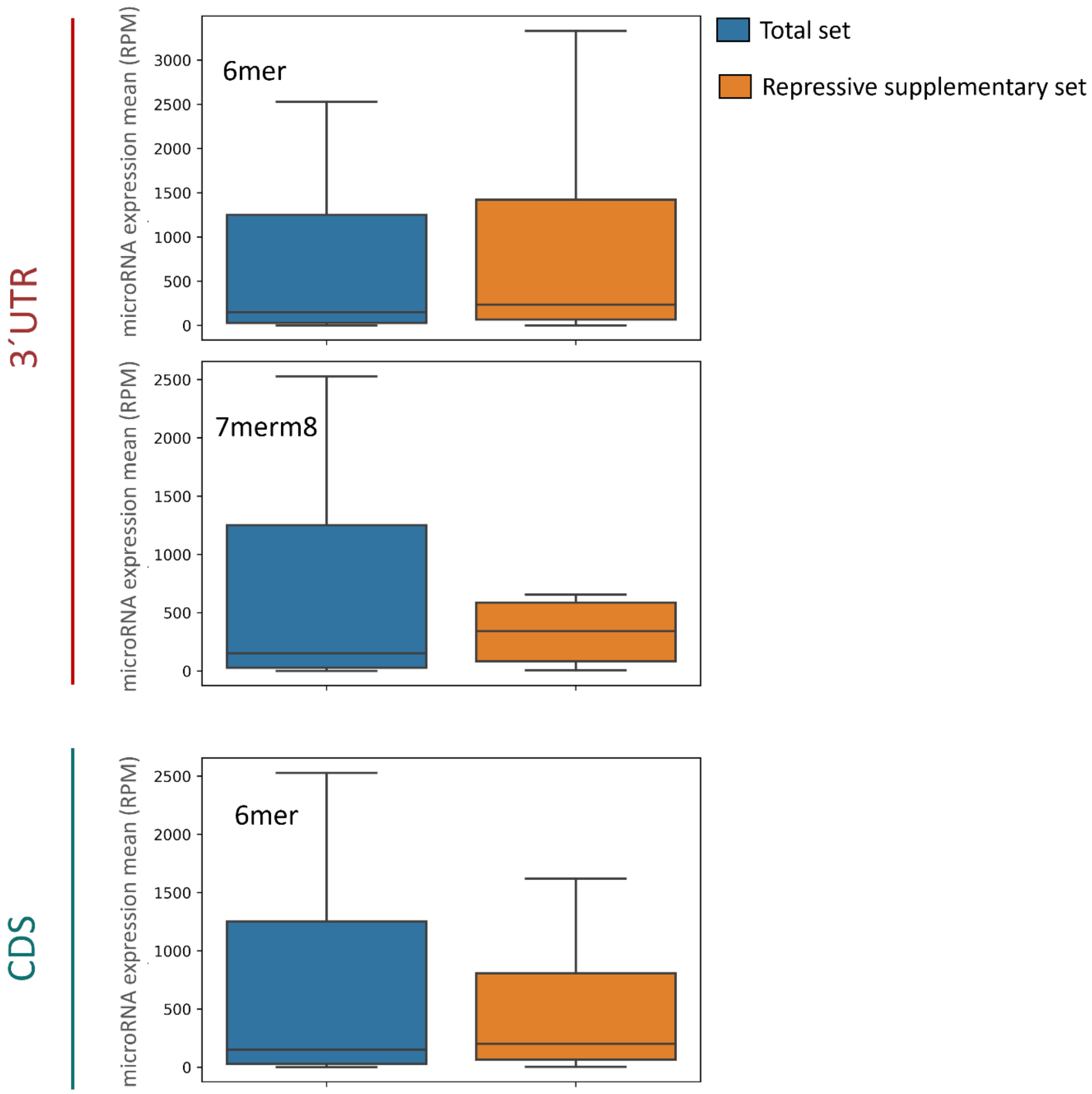


**Supplementary Figure 4.** *Mean microRNA expression of the 143 microRNAs studied (blue) and the 39 microRNAs with repressive supplementary interactions (orange).* The interactions at the indicated transcript regions are plotted separately. Only the regions that showed a statistically significant contribution of the 3´-supplementary interactions were included in the analysis (19: 6mer at the 3´UTR, 16: 7mer-m8 at the 3´UTR and 21: 6mer at the CDS).

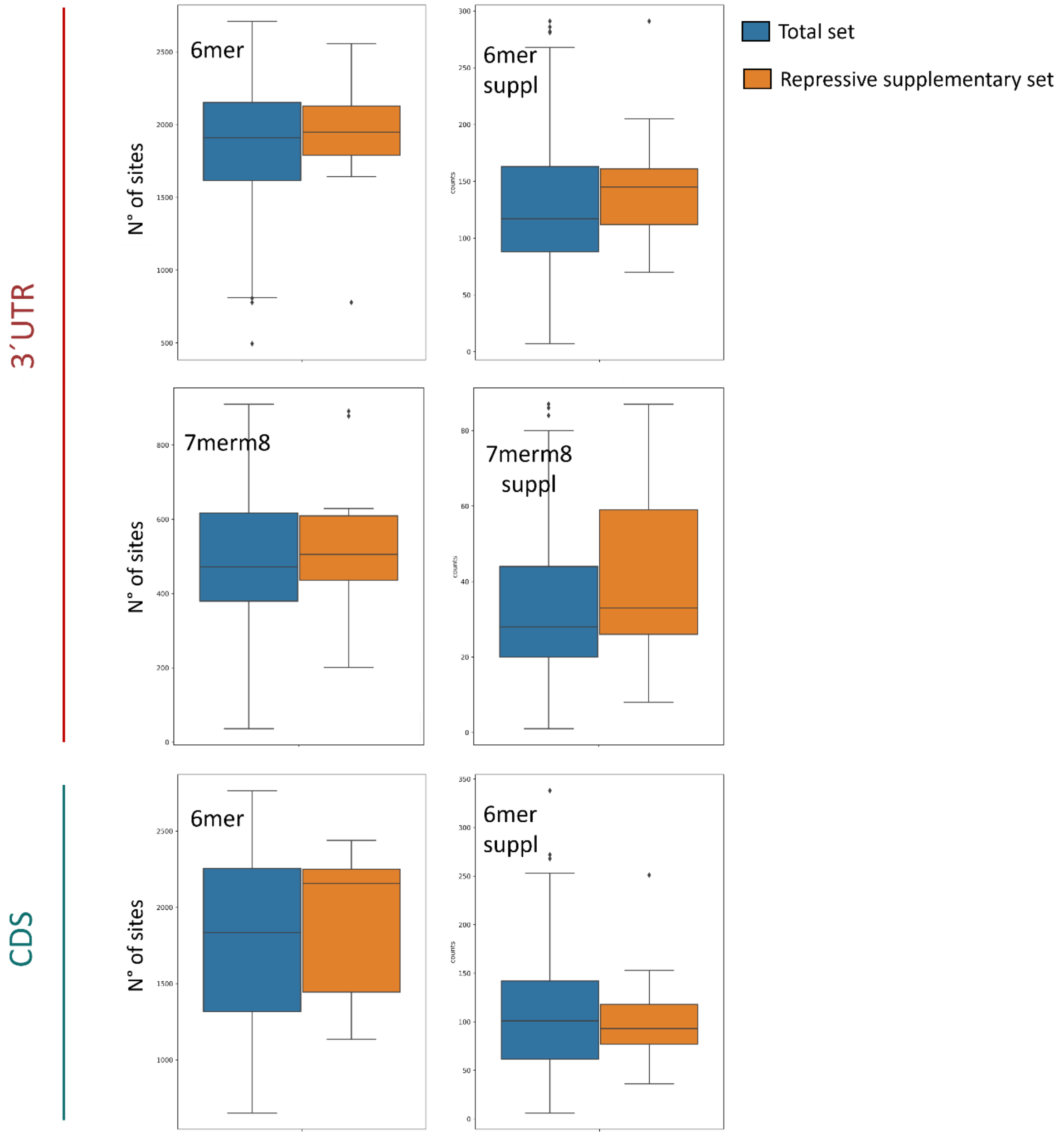


**Supplementary Figure 5.** Number of sites for the 143 microRNAs studied (blue) and the 39 microRNAs with repressive supplementary interactions (orange). The interactions at the indicated transcript regions are plotted separately. Only the regions that showed a statistically significant contribution of the 3´-supplementary interactions were included in the analysis (19: 6mer at the 3´UTR, 16: 7mer-m8 at the 3´UTR and 21: 6mer at the CDS).

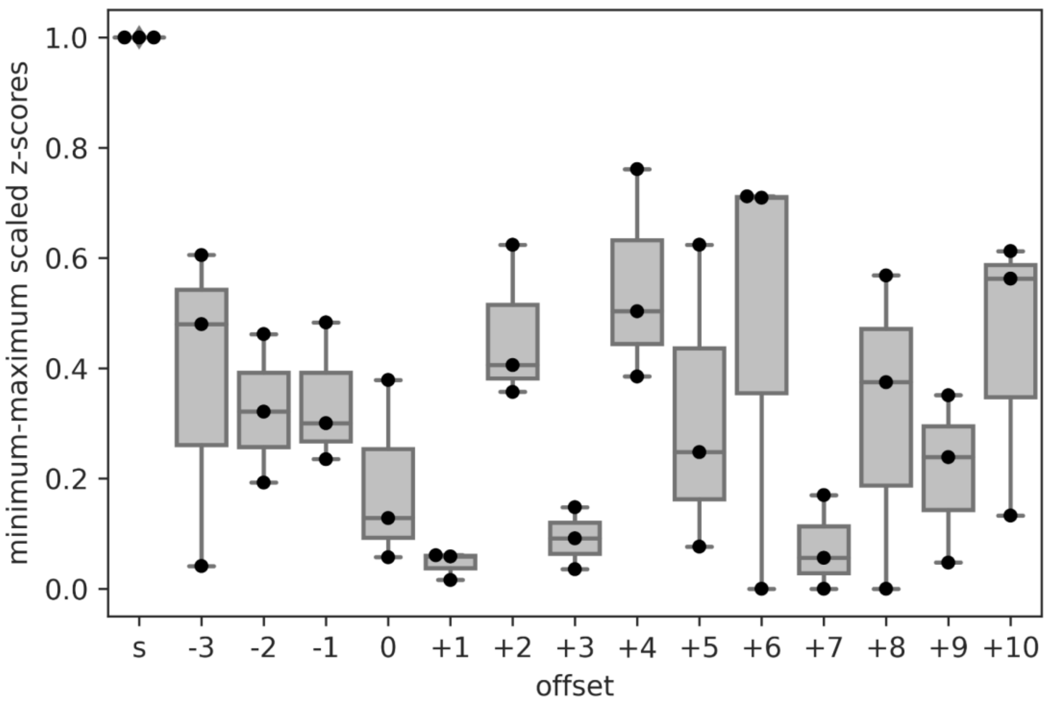


**Supplementary Figure 6.** *Distribution of scaled Z-scores by offset of the microRNA/mRNA interaction using 3'-supplementary pairing region.* Each boxplot is composed by 3 datapoints which correspond to the Z-score mean of each interaction (6mer 3'UTR, 7mer-m8 3'UTR, 6mer CDS). Data was minimum-maximum scaled inside each interaction to normalize the different ranges of Z-scores and then we combine the three median of the three Z-scores.

### Supplementary material - Table List

**Supplementary_Table_1_Studied_transcripts_microRNA_and_mRNAs**

**Supplementary_Table_2_Counts_microRNA-mRNA_interactions**

**Supplementary_Table_3_FDR_seed_suppl_comparisons**

**Supplementary_Table_4_individualMicroRNAstats**
